# Porcine sapovirus protease controls the innate immune response and targets TBK1

**DOI:** 10.1101/2023.12.21.572846

**Authors:** Iliana Georgana, Myra Hosmillo, Aminu S. Jahun, Edward Emmott, Frederic Sorgeloos, Kyoung-Oh Cho, Ian G. Goodfellow

## Abstract

Human sapoviruses (HuSaVs) and noroviruses are considered the leading cause of acute gastroenteritis worldwide. While extensive research has focused on noroviruses, our understanding of sapoviruses (SaVs) and their interactions with the host’s immune response remains limited. HuSaVs have been challenging to propagate in vitro, making the porcine sapovirus (PSaV) Cowden strain a valuable model for studying SaV pathogenesis. In this study we show, for the first time, that PSaV Cowden strain has mechanisms to evade the host’s innate immune response. The virus 3C-like protease (NS6) inhibits type I IFN production by targeting TBK1. Catalytically active NS6, both during ectopic expression and during PSaV infection, targets TBK1 which is then led for rapid degradation by the proteasome. Moreover, deletion of TBK1 from porcine cells led to a significant increase in PSaV titres, emphasizing its role in regulating PSaV infection. Additionally, we successfully established PSaV infection in IPEC-J2 cells, an enterocytic cell line originating from the jejunum of a neonatal piglet. Overall, this study provides novel insights into PSaV evasion strategies, opening the way for future investigations for SaV-host interactions, and enabling the use of a new cell line model for PSaV research.

## 1. Introduction

Sapoviruses (SaVs) are small non-enveloped RNA viruses belonging to the *Caliciviridae* family. Together with noroviruses they are the main cause of acute gastroenteritis worldwide in both children and adults. SaVs are classified into at least 19 genogroups (GI to GXIX) [1,2], depending on their genetic diversity, and they have been found to infect various animal species and humans. SaVs have a positive-sense, single-stranded RNA genome ranging from 7.1 to 7.7 kbp, which is linked at the 5’ end to the virus-encoded VPg protein. The SaV genome contains two open reading frames (ORFs), ORF1 and ORF2. An additional ORF (ORF3) has been predicted in several sapovirus strains, however its function is still unknown [3]. ORF1 encodes all non-structural proteins and the major structural protein VP1 [4], ORF2 encodes the minor structural protein VP2 [5]. ORF1 is translated to a polyprotein that undergoes proteolytic processing by the virus encoded 3C-like protease NS6. In SaVs, the protease is found as a protease-polymerase fusion protein and not further processed into the mature NS6 protease and NS7 RNA polymerase domains [6]. This characteristic shared with feline caliciviruses (FCV) [7], but not observed in any other *Caliciviridae* member.

Among the SaV strains infecting humans, genogroups GI and GII are the primary cause of gastroenteritis outbreaks as well as sporadic cases. For over 40 years since its discovery [8,9], human sapovirus (HuSaV) could not be propagated *in vitro* until recently [10-12]. Prior to these advancements, the porcine sapovirus (PSaV) Cowden strain (belonging to genogroup GIII) served as a valuable model for studying SaV pathogenesis and it remains the only SaV for which a robust reverse genetics system is available. The ability of this strain to replicate in porcine kidney cells *in vitro* [13] and the establishment of a reverse genetics system [14] made it an appealing model with which to uncover the basic mechanisms of SaV biology.

PSaV is most likely endemic in pigs worldwide. It has been detected in fecal samples of domestic pigs from around the world, and in wild boars in Japan [15]. Although so far there is no reported SaV transmission between swine and human [15], there were concerns for potential zoonotic infections or recombination events, due to high genetic similarity of some PSaV strains and HuSaV [16]. While there is extensive research on the interplay between noroviruses and the host immune response [17], our understanding of SaV in this context is limited. Nevertheless, it is known that SaVs exhibit sensitivity to type I IFN [18], indicating the involvement of innate immune pathways in controlling SaV infections, and highlighting the need to understand the potential mechanisms employed by SaVs to evade the host’s immune response.

Innate immunity is the primary line of defense against viruses and other pathogens. In part, it relies on pattern recognition receptors (PRRs), such as retinoic acid-inducible gene-I (RIG-I)-like receptors (RLRs) [19], which detect pathogen-associated molecular patterns (PAMPs). RIG-I and melanoma differentiation associated gene 5 (MDA5) are RLRs that recognise viral RNA genome and RNA replication intermediates. Recognition of these PAMPs triggers a cascade of signalling events via the activation of mitochondrial antiviral-signalling protein (MAVS) [19,20]. MAVS signals via the TANK-binding kinase 1 (TBK1) and Inhibitor of nuclear factor-κB Kinase ε (IKKε) proteins, leading to the activation of IRF3 and IRF7 transcriptional factors [21]. Upon their activation, these transcription factors translocate to the nucleus and induce transcription of type I interferons (IFNs) and other antiviral or immunoregulatory genes [20,22].

Here we show that PSaV Cowden strain has mechanisms to counteract the host innate immune response. Specifically, we have found that the virus protease (NS6) inhibits the type I IFN response by targeting TBK1 for proteasomal degradation in a manner dependent on its protease activity. TBK1 protein levels are significantly decreased during PSaV infection and they are completely rescued when the proteasome was inhibited. Depletion of TBK1 from cells, led to a significant increase of infectious PSaV titres. This highlights the critical role of TBK1 in regulating PSaV infection, and elucidates the rationale behind the targeting of TBK1 by NS6. Finally, here for the first time we established PSaV infection in IPEC-J2 cells, enteric epithelial cells, originally derived from the jejunum of a neonatal piglet [23]. In conclusion, our study reveals that PSaV Cowden strain, the only PSaV strain adapted to grow *in vitro*, possesses mechanisms to evade the host’s innate immune response. Our research shows that SaVs have mechanisms to counteract their host’s response, and paves the way for further investigations into SaV-host interactions.

## 2. Materials and Methods

### 2.1 Cell lines and virus

HEK293T (ATCC CRL-11268), were grown in Dulbecco modified Eagle’s medium (DMEM, Sigma Aldrich), Parental LLC-PK1 (ATCC CL-101) cells and LLC-PK1 EV or KO (described in 2.7) were grow in Minimal Essential Medium (MEM, Gibco). For all cell lines, media were supplemented with 10% (v/v) heat-inactivated Fetal Bovine Serum (FBS) (HyClone), 10 U/ml of penicillin, 100 mg/mL of streptomycin, 2 mM L-glutamine (Sigma Aldrich), and non-essential amino acids (Sigma Aldrich). IPEC-J2 (DSMZ ACC 701) cells were cultured in Advanced Dulbecco’s Modified Eagle Medium/Ham’s F12 (Advanced DMEM/F12, Gibco), supplemented with 10% heat-inactivated FBS (HyClone), 10 U/ml of penicillin, 100 mg/mL of streptomycin, 2 mM L-glutamine (Sigma Aldrich), 15 mM HEPES (Invitrogen) and non-essential amino acids (Sigma Aldrich). The maintenance of IPEC-J2 cells was adapted from [24]. All cell lines were grown at 37 °C with 5% CO2. Infectious particles of a tissue-culture adapted PSaV Cowden strain were recovered from full-length infectious cDNA clone pCV4A (a kind gift from Dr K. O. Chang, Kansas State University, Manhattan, KS, USA). Virus was propagated in LLC-PK1-NPro cells [18] and purified by sedimentation through a cushion of 30% (w/v) sucrose.

### 2.2 Expression vectors

Tissue culture-adapted porcine enteric calicivirus sequence in the GenBank database (accession number AF182760) was used to generate protease-polymerase (Pro^WT^Pol^WT^) fusion constructs into an empty vector (InvivoGen). Mutation of the active site in either protease, polymerase, or both, were made to produce Pro^MUT^Pol^WT^, Pro^WT^Pol^MUT^ and Pro^MUT^Pol^MUT^, respectively. In the protease the Cys of the active site was mutated into Ala, and in the polymerase the active site Gly-Asp-Asp was mutated into an Gly-Ala-Ala. The Pro-Pol fusion HA-tagged (Pro^WT^Pol^WT^ and Pro^MUT^Pol^WT^) plasmids were generated using an in-house human PGK promoter-driven lentiviral vector via Gateway cloning (Invitrogen). Plasmids expressing HuSaV (GI.2) and PSaV protease-polymerase fusion were generated by cloning into the XhoI and BamHI sites of pmCherry-C1 (Clontech). Vectors expressing RIG-I-CARD domain [25] and IRF3-5D [26] were previously described. Full length RIG-I (pUNO-hRIG-I) used in the 5Br assay was obtained from InvivoGen. To generate the FLAG-pTBK1 plasmid, full length porcine TBK1 was first synthesized by PCR using cDNA from LLC-PK1 cells as template (forward primer: 5’-AAAAACCGGTACCATGCAGAGCACTTC-3’, reverse primer: 5’-AAAACGTACGACTAAAGACAGTCAAC-3’), digested with AgeI and BsiWI, and then ligated into an in-house human PGK promoter-driven lentiviral vector in frame with 3 tandem N-terminal FLAG tags. The FLAG-pTBK1-HA plasmid was generated by Gateway cloning (Invitrogen), using the FLAG-pTBK1 as template for generation of the entry clone (forward primer: 5’-GGGGACAAGTTTGTACAAAAAAGCAGGCTTCACCATGGATTACAAGGATGACGAC-3’, reverse primer: 5’-GGGGACCACTTTGTACAAGAAAGCTGGGTTAAGACAGTCAACATTGCG-3’), and an in-house human PGK promoter-driven expression clone containing a gateway cassette in frame with 3 tandem C-terminal HA tags. The GFP-FLAG plasmid was a gift from Prof. Geoffrey L. Smith (University of Oxford). The IFN-β firefly luciferase reporter plasmid was from T. Taniguchi (University of Tokyo, Japan), and pTK-renilla luciferase plasmid was from Promega.

### 2.3 Transfection of cell lines

HEK293T cells were transfected with polyethylenimine (PEI, CellnTec, 3 μL per μg of DNA for 96-wp or 6-wp, and 5 μL per μg of DNA for 10 cm dishes), unless indicated otherwise. LLC-PK1 cells were transfected with Lipofectamine 2000 (Invitrogen), according to the manufacturer’s instructions.

### 2.4 Reporter gene assays

HEK293T were seeded in 96-wp and 24 h later cells were transfected with 70 ng per well of the firefly luciferase reporter plasmids (IFN-β), together with 10 ng per well of the pTK-Renilla luciferase plasmid and the plasmids encoding the genes of interest, or empty vector (EV). The exact amounts of expression plasmids used are included in the figure legends. To ensure consistency, EV was added to the transfection when needed, maintaining a constant final amount of transfected DNA. In instances where stimulation was carried out by transfecting another plasmid, an equivalent amount of EV was transfected to the non-stimulated (NS) control wells. Transfection was performed with PEI (CellnTec, 3 μL per 1 μg DNA) according to the manufacturer’s instructions. DMEM supplemented with 2% (v/v) FBS was then added to achieve a volume of 100 μL per well. Cells were incubated at 37 °C for 24 h and then they were washed with 200 μL PBS per well and harvested in 100 μL per well of passive lysis buffer (Promega). Cells were freeze-thawed and 25 μL of lysate from each well was added to 25 μL of firefly luciferase reagent (Bright-Glo, Promega), and 10 μL of each lysate was added to 50 μL of 2 μg/mL Renilla luciferase substrate (NanoLight Technology). Measurement of luciferase activity was carried out using a Glomax Navigator Microplate Luminometer (Promega). In all experiments, cells were transfected in triplicate. Each firefly-luciferase reading was normalised to its corresponding renilla-luciferase reading. To calculate fold inductions, these normalised values were divided by the non-stimulated control value for each plasmid.

### 2.5 5Br assay

The 5Br assay was conducted as previously described [27]. Briefly, HEK293T cells were seeded in 96-wp and 24 h later they were transfected with 20 ng of IFN-β reporter, 5 ng of pTK-Renilla reporter, 0.5 ng of pUNO-hRIG-I, and 50 ng of any of the plasmids expressing Pro^WT^Pol^WT^, Pro^WT^Pol^MUT^, Pro^MUT^Pol^MUT^ and Pro^MUT^Pol^WT^. Transfections were conducted with Lipofectamine 2000 (Invitrogen) according to the manufacturer’s instructions. Where necessary, EV was used to maintain a constant amount of plasmid DNA per transfection. Four h post-transfection, medium was changed with fresh medium containing 2 μg/mL of poly(I:C). A non-poly(I:C) control medium was included as non-stimulated (NS) condition. Luciferase activity was measured 36 h post-transfection using the Dual-Glo luciferase assay system (Promega) by using the Glomax Navigator Microplate Luminometer (Promega). The ratios of firefly luciferase to Renilla luciferase activities were calculated for each sample, and these values were then divided by the non-stimulated EV control value to calculate fold change.

### 2.6 Poly(I:C) stimulation and RT-qPCR

TBK1 WT (EV) and KO LLC-PK1 cells were seeded in 12-wp and 24 h later they were transfected with 1 μg/mL poly(I:C) (P1530, Sigma) with 3 μL per well of Lipofectamine 2000 (Invitrogen). A non-transfected condition was included as non-stimulated (NS) control. Six h later, cells were lysed in 250 μL per well of RNA lysis buffer (4 M guanidine thiocyanate, 25 mM Tris pH 7). Total RNA extraction with an on-column DNAse treatment was performed with the GenElute Mammalian Total RNA Miniprep Kit (Sigma Aldrich) according to the manufacturer’s protocol. Five μL of each RNA sample were used for cDNA synthesis, which was performed with the M-MLV Reverse Transcriptase (Promega), according to the manufacturer’s instructions. qPCR was carried out using a 2X SYBR Green mastermix containing 2.5 mM MgCl2, 400 mM dNTPs, 1/10,000 SybrGreen (Molecular Probes), 1 M Betaine (Sigma), 0.05 U/mL of Gold Star polymerase (Eurogentec), 1/5 10X Reaction buffer (750 mM Tris-HCl pH 8.8, 200 mM [NH4]2SO4, 0.1% [v/v] Tween 20, Without MgCl2), and ROX Passive Reference buffer (Eurogentec), and ran on a ViiA 7 Real-Time PCR System (ThermoFisher Scientific), with a 15 sec 95 °C denaturation step and a 1 min 60 °C annealing/extension step for 40 cycles. The primers used were: porcine *IFN-β* (Fwd: 5’-GGAGCAGCAATTTGGCATGT-3’; Rv: 5’-TGACGGTTTCATTCCAGCCA-3’) and porcine *β-actin* (Fwd: 5’-TCTACACCGCTACCAGTTCG-3’; Rv: 5’-GCTCGATGGGGTACTTGAGG-3’). Gene amplification was normalised to *β-actin* housekeeping gene and the fold induction was calculated relative to the unstimulated EV sample.

### 2.7 CRISPR/Cas9-mediated genome editing

Guide RNA (gRNA) for the porcine TBK1 was designed using an online software (https://portals.broadinstitute.org/gpp/public/analysis-tools/sgrna-design) to target exon 3, which is shared by 5 out of 6 isoforms. The 6^th^ isoform is predicted to give rise to a protein of 64 kDa size, which has not been identified in our study. The gRNA complex used in this study is: 5’-CACCGACAGTGTATAAACTCCCACA-3’ and 5’-AAACTGTGGGAG-TTTATACACTGTC-3’. CRISPR/Cas9-mediated genome editing was performed as described previously [28]. Briefly, the oligo complexes were annealed, phosphorylated and then ligated into the PX459 CRISPR/Cas9 vector. Plasmid with or without the gRNA sequence was transfected into LLC-PK1 cells with Lipofectamine 2000 (Invitrogen) and cells were incubated for 48 h. Then puromycin (4 μg/mL) was added for 24 h and puromycin-resistant cells were seeded in 96-wp at a density of 0.5 cell/well and were left to form colonies for 2-3 weeks. Several clones were expanded, and potential knockout clones were selected by immunoblotting and confirmed by DNA sequencing at the gRNA target site. Primers used to amplify and sequence *TBK1* exon 3 are: Fwd: 5’-TTTAACAAACCCCAGGTAAA-3’ and Rv: 5’-ACCTGGCTTGGATCCATGGT-3’. In the TBK1 KO cell line both alleles were observed to have the same frameshifting, resulting in no full length TBK1 production.

### 2.8 Immunoprecipitation (IP)

HEK293T cells were seeded in 10-cm dishes and transfected with the indicated plasmids and amounts as described in the figure legend. After 24 h, cells were washed once with PBS and then lysed with 0.5% NP-40 in PBS supplemented with cOmplete Mini EDTA-free protease inhibitor cocktail (Roche) and Benzonase (Merck Millipore, 1 μL per mL of lysis buffer). Lysates were incubated at 4 °C on an end-to-end rotor for 20 min, and then insoluble material was removed by centrifugation at 21,000 × *g* for 20 min at 4 °C and discarded. Lysates were then incubated with Anti-FLAG M2 Affinity Gel (Sigma Aldrich) agarose gel beads (30 μL per sample) at 4 °C on an end-to-end rotor for 4 h. Subsequently, the beads were washed 3 times with the lysis buffer, and finally were incubated at 100 °C for 5 min in loading buffer to elute bound proteins. Cleared lysates and IP eluates were analysed by SDS-PAGE and immunoblotting. Data shown are representative of at least 2 independent experiments showing similar results.

### 2.9 SDS-PAGE and immunoblotting

Cells were lysed in 0.5% NP-40 in PBS for the IPs, or in RIPA buffer (50 mM Tris pH 8, 150 mM NaCl, 1 mM EDTA, 1% Triton X-100, 0.1% SDS) for all the other experiments. Both lysis buffers were supplemented with cOmplete Mini EDTA-free protease inhibitor cocktail (Roche) and Benzonase (Merck Millipore, 1 μL per mL of lysis buffer). Cells in 6-wp and 10 cm dishes were lysed in 200 μL and 500 μL of lysis buffer, respectively. Lysates were incubated on an end-to-end rotor for 20 min at 4 °C and then spun down at 21,000 × *g* for another 20 min at 4 °C. Supernatants were transferred to a new tube and the pellets were discarded. Samples were quantified using the BCA assay (Thermo Scientific) according to the manufacturer’s recommendations and made up to the desired concentration with lysis buffer. The samples were then mixed with loading buffer (0.6% SDS, 8% glycerol, 0.002% bromophenol blue, 25 mM Tris pH 6.8, 0.02% 2-mercaptoethanol), heated at 100 °C for 5-20 min and then were resolved by SDS-PAGE or NuPAGE (see each figure legend). Proteins were transferred to 0.45 μm nitrocellulose membranes (GVS) using a Trans-Blot semi-dry transfer unit (Bio-Rad), for 40 mins at 25 V. Membranes were blocked in 0.1% Tween PBS supplemented with 5% (w/v) skimmed milk and subjected to immunoblotting with the following primary antibodies at the indicated dilutions: TBK1 (Cell Signaling [CS-3013], rabbit polyclonal, 1:500); GAPDH (Invitrogen [AM4300], mouse monoclonal, 1:5,000); RIG-I (Santa Cruz [sc-376845], mouse monoclonal, 1:500); FLAG (Sigma [F3165], mouse monoclonal, 1:1,000); HA (Abmart [M20003M], mouse monoclonal, 1:5,000); and mCherry (Cell Signaling [CS-43590], rabbit monoclonal, 1:1,000). The PSaV VPg and NS6 antibodies were generated by immunisation of New Zealand White rabbits with purified recombinant VPg and NS6, respectively. Primary antibodies were detected using IRDye-conjugated secondary antibodies in an Odyssey Infrared Imager (LI-COR Biosciences). All secondary antibodies were used at 1:10,000 dilution.

### 2.10 PSaV infection

For detecting TBK1 levels during infection, LLC-PK1 cells in 6-wp were infected with PSaV at 5 TCID50/cell supplemented with 200 μM GCDCA. Incubation with inocula was for 1 h at room temperature (RT) on a shaker. Unabsorbed virus was then removed and cells were washed once with MEM. Fresh MEM supplemented with 200 μM GCDCA was then added on the cells in the presence or absence of Z-VAD-FMK (Promega), NH4Cl (Fisher Scientific), MG132 (Selleck chemicals), lactacystin (Cayman), or DMSO (Sigma). Cells were incubated at 37 °C for the time post-infection (pi) indicated in each figure legend. For measuring virus yield in TBK1 WT and KO LLC-PK1 cells, cells were infected in suspension with PSaV at 1 TCID50/cell supplemented with 200 μM GCDCA. Incubation with inocula took place for 1 h at 37 °C on an end-to-end rotor. Cells were then washed once, resuspended in fresh MEM and seeded on 12-wp. Total virus (cells and supernatants) was then used for TCID50 analysis. For infection of IPEC-J2 cells, cells in 6-wp were infected with PSaV at 5 TCID50/cell with or without 300 μM GCDCA. Incubation with inocula was for 3 h at RT on a shaker. Unabsorbed virus was washed and fresh MEM supplemented with 300 μM GCDCA was added on the cells.

### 2.11 TCID50 assay

All TCID50s were performed in parental LLC-PK1 cells, seeded in 96-wp. Ten-fold serial dilutions of clarified virus supernatants were prepared in MEM supplemented with GCDCA (final concentration 200 μM) and aliquoted into wells of 96-wp, each in 4 replicates of 50 μL. Seven days pi, cells were assessed for cytopathic effect (CPE), and TCID50/mL was calculated by the Reed–Muench method [29].

### 2.12 Statistical analysis and software

Statistical significance was determined using an unpaired Student’s *t*-test by using GraphPad Prism 9 software. All experiments were carried out at least twice, and one representative data set of each experiment is shown. Statistical analysis was performed within a single experiment. In all cases, ns: non-significant, \**p* ≤ 0.05, \*\**p* ≤ 0.01, \*\**\*p* ≤ 0.001, and \*\*\*\**p* ≤ 0.0001. Image Studio Lite 5.2 was used for Western blot quantification. Clustal Omega was used for all sequence alignments, and Snapgene 4.2 was used for primer design and cloning strategies. The schematic on Figure 1A was created on BioRender.com. ChatGPT (GPT-3.5) was used for editing and proofreading.

**Figure 1.**
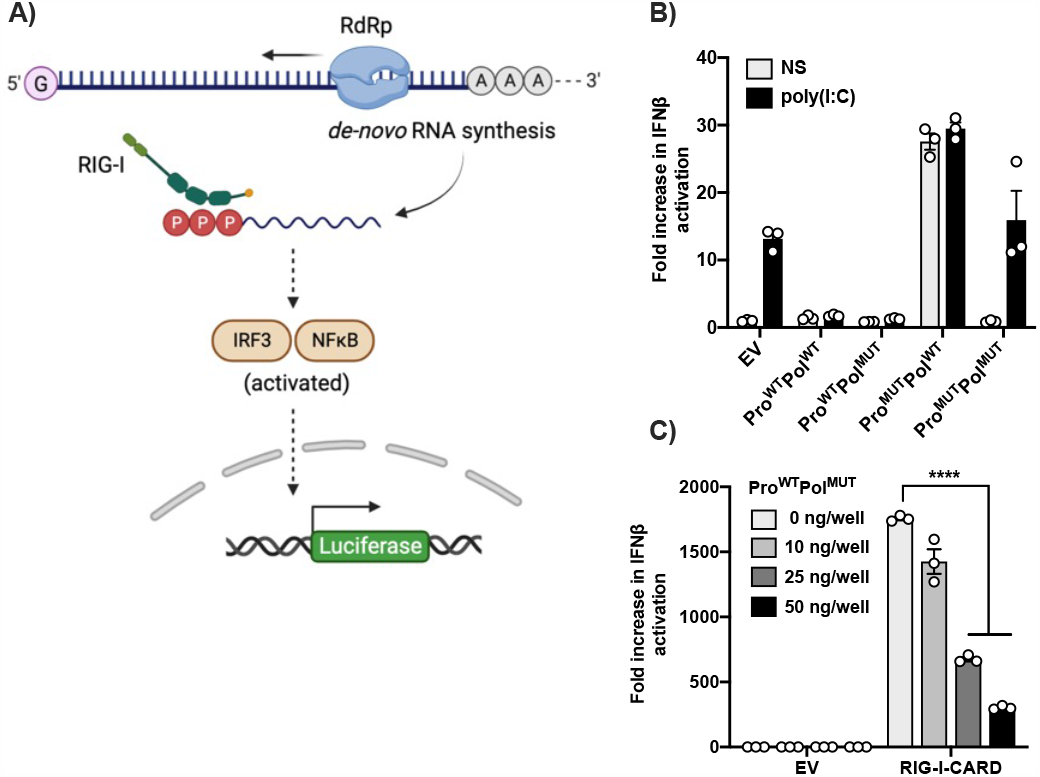
PSaV NS6 is a type I IFN antagonist. A) Schematic of 5Br assay, first described in [27]. B) HEK293T cells were transfected in triplicate with a reporter plasmid encoding firefly luciferase under the control of IFN-β promoter, a renilla luciferase transfection control, a RIG-I expressing plasmid and 50 ng/well of any of the indicated Pro-Pol fusion plasmids. Four h post-transfection, cells were stimulated with poly(I:C) (2 μg/mL), or left unstimulated (NS). Cells were harvested in passive lysis buffer 36 h post-transfection and firefly luciferase activity was measured and normalised to the renilla luciferase activity. Each sample was compared to the average value of EV control to obtain the fold increase. C) HEK293T cells were transfected in triplicate with the IFN-β reporter, the renilla luciferase control reporter, and the indicated amounts of Pro^WT^-Pol^MUT^ fusion construct. Empty vector (EV) was added to samples when necessary to keep the final amount of DNA transfected as 50 ng/well in all samples. Stimulation was achieved by co-transfection of RIG-I-CARD expression plasmid (5 ng/well). After 24 h, cells were lysed in passive lysis buffer, and firefly luciferase was measured and normalised to the renilla luciferase. Fold change was obtained by comparing each stimulated sample to the average value of its unstimulated control. Data shown are representative of 3 independent experiments where each independent experiment was performed with a minimal of 3 wells for each condition. Statistical significance was measured by a Student’s *t-*test. \*\*\*\**p* ≤ 0.0001. Error bars represent standard error of the mean.

## 3. Results

### 3.1 PSaV NS6 is a type I IFN antagonist

PSaV is sensitive to IFN response [18], and therefore we explored potential counteractive mechanisms that it might possess. There is a well-established precedent for a role of viral proteases in the regulation of the IFN response [30]. Recent studies on the human norovirus protease have indicated its capability to cleave NEMO, the regulatory subunit of the IKK complex [31], although it is unclear how this cleavage impacts the viral life cycle. In SaVs the protease (NS6) is produced as a fusion with the polymerase (NS7). RNA-dependent RNA polymerases (RdRps), when ectopically expressed, have been shown to generate uncapped 5’-triphosphate RNAs, which are recognized by RIG-I [32,33], therefore driving a robust type I IFN response. This capability can be used to establish a reporter readout in cells that relies on the activity of the authentic viral polymerase and has previously been described, the 5Br assay [27]. Briefly, over-expression of an RdRp, NS7 in the case of noroviruses, induces downstream activation of signalling pathways, such as IRF3 and NF-κB. Co-transfection of a luciferase reporter where the luciferase gene is under the control of the type I IFN promoter enables the measurement of this activation (Figure 1A). To determine if NS6 is an IFN antagonist, we examined the ability of NS6 to inhibit the response driven by NS7 over-expression in this assay. We generated constructs encoding the PSaV NS6-7 fusion where one of the two, or both, were catalytically active, or inactive. We co-transfected these constructs, together with an IFN-β reporter, and we further stimulated the cells with poly(I:C) transfection or left them unstimulated (NS). Active NS7 (Pol^WT^) induced a robust IFN-β activation, but this was only observed when NS6 was catalytically inactive (Pro^MUT^). Active NS6 (Pro^WT^) significantly inhibited IFN-β activation driven by either NS7 RdRp activity, or by poly(I:C) transfection (Figure 1B), indicating that it is a type I IFN antagonist. To further demonstrate that this effect was driven through RIG-I mediated sensing, we employed reporter gene assays where IFN-β activation was achieved by ectopic expression of RIG-I CARD domains. The IFN-β reporter and the RIG-I-CARD domains were co-expressed with increasing doses of NS6-7, where NS7 was catalytically inactive to avoid background reporter activation. NS6 restricted IFN-β activation in a dose-response and statistically significant manner (Figure 1C). Overall, these results show that NS6 strongly inhibits type I IFN activation.

### 3.2 NS6 restricts the type I IFN pathway at the level of TBK1

To further understand how NS6 antagonizes the IFN response, we sought to investigate where on the pathway the inhibitory function is exerted. For this, we again employed reporter gene assays and we stimulated IFN by over-expression of components of the pathway, such as RIG-I-CARD, TBK1 and the constitutively active IRF3-5D [26]. Ectopic expression of these proteins results in activation of the pathway at that particular point and its subsequent activation downstream. These molecules were co-expressed together with the IFN-β reporter and the NS6-7 (Pro-Pol) fusion, where NS6 was catalytically active (WT), or inactive (MUT). In both cases, NS7 was catalytically inactive (MUT), to prevent signal transduction caused by the *de novo* initiation activity. We observed that only the expression of a catalytically active NS6 significantly reduced IFN-β activation downstream of both RIG-I and TBK1 over-expression (Figure 2A-B). Inhibition was not observed when activation was driven by overexpression of IRF3 (Figure 2C), suggesting that NS6 acts at the level of TBK1.

**Figure 2.**
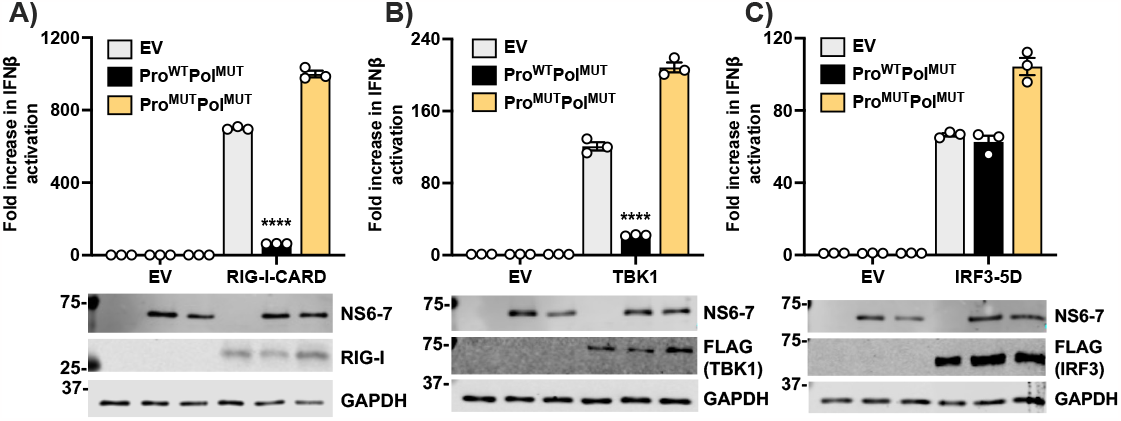
PSaV NS6 restricts type I IFN activation at the level of TBK1. HEK293T cells were transfected in triplicate with the IFN-β firefly luciferase reporter, the renilla luciferase control reporter, and 50 ng/well of EV, Pro^WT^-Pol^MUT^ or Pro^MUT^-Pol^MUT^. Cells were also co-transfected with EV as a non-stimulated control, or A) RIG-I-CARD (5 ng/well), B) TBK1 (40 ng/well), or C) IRF3-5D (40 ng/well) expression plasmids to activate the type I IFN pathway. Twenty-four h post-transfection, cells were lysed, and firefly luciferase was measured and normalised to the renilla luciferase. Each stimulated sample was compared to the average value of its unstimulated control, to obtain fold increase. Immunoblots underneath each graph show expression levels of the different overexpressed proteins, or GAPDH. The positions of molecular mass markers in kDa are indicated on the left of each immunoblot. Data shown are representative of 3 independent experiments, where each independent experiment was performed with a minimal of 3 wells for each condition. Statistical analysis was performed within a single experiment, and significance was measured by a Student’s *t-*test. \*\*\*\**p* ≤ 0.0001. Error bars represent standard error of the mean.

### 3.3 NS6 interacts with and modulates the protein levels of TBK1

Our findings so far demonstrate that NS6 restricts type I IFN response at the level of TBK1 (Figures 2). To further investigate if NS6 specifically targets TBK1 for cleavage, we co-expressed HA-tagged NS6-7 fusion proteins, where NS6 is active (WT), or inactive (MUT), with FLAG-tagged porcine TBK1 (pTBK1), or GFP as a control. Although no cleavage product was observed, the total level of TBK1 protein was reduced in the presence of WT NS6, but not in the presence of a catalytically inactive NS6, while the level of GFP remained unaffected (Figure 3A). To investigate this further and determine if this was due to limitations of the TBK1 antibody to detect a cleaved product, we used a double-tagged pTBK1 construct. We cloned pTBK1 into a mammalian expression vector with a FLAG-tag on its N-terminus and an HA-tag on its C-terminus. We overexpressed the double-tagged construct with catalytically active or inactive NS6 alongside the single-tagged pTBK1 (FLAG-pTBK1) as a control. No cleavage product was detected after immunoblotting with either anti-FLAG or anti-HA antibodies (Figure 3B). Nevertheless, the reduction of TBK1 protein levels was consistent and independent of whether TBK1 was single or double tagged.

**Figure 3.**
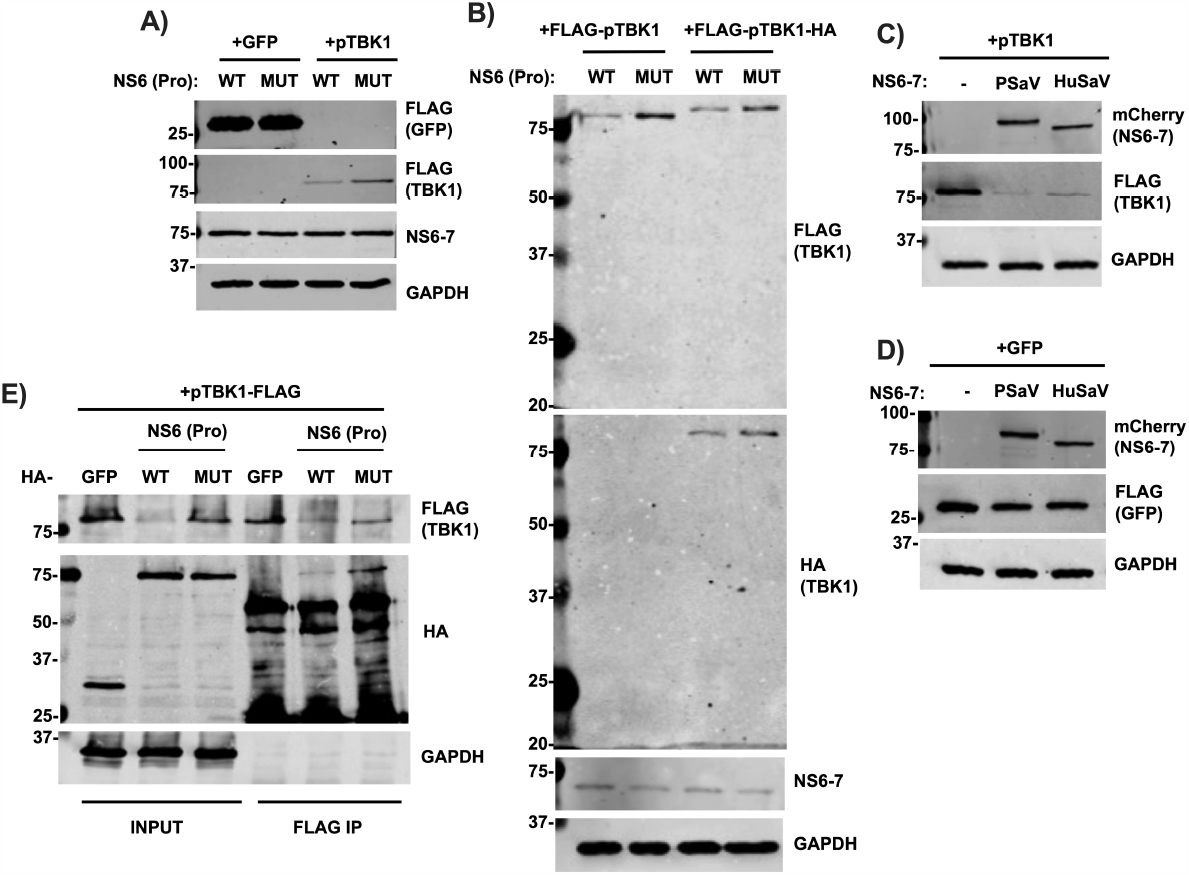
NS6 interacts with TBK1 and modulates its protein levels. A) HEK293T cells were transfected for 24 h with 2 ug of HA-NS6-7 where NS6 was WT (Pro^WT^Pol^WT^), or MUT (Pro^MUT^Pol^WT^), together with 2 ug of FLAG-pTBK1 or 0.5 ug of GFP-FLAG. Cells were lysed, loaded onto an SDS-PAGE (2 μg/sample), separated and analysed by immunoblotting with the indicated antibodies. B) HEK293T cells were co-transfected with 2 ug HA-tagged NS6-7 WT or MUT (as described above), together with 2 ug of FLAG-pTBK1, or FLAG-pTBK1-HA. Cells were lysed 24 h later, and samples were loaded onto an SDS-PAGE (2 μg/sample), separated and immunoblotted with the indicated antibodies. C-D) HEK293T cells were transfected with 2 ug of EV, or PSaV or HuSaV NS6-7, both mCherry-tagged, along with B) 2 ug of FLAG-pTBK1, or C) 0.5 ug GFP-FLAG. Twenty-four h later, cells were lysed, loaded onto an SDS-PAGE (2 μg/sample), separated and analysed by immunoblotting with the indicated antibodies. E) HEK293T cells were co-transfected with 4 ug FLAG-pTBK1 and 4 ug of HA-Pro^WT^Pol^WT^, or HA-Pro^MUT^Pol^WT^ (also mentioned HA-NS6-7 in the text), or 1 ug HA-GFP. Cells were harvested 24 h later, lysed, and subjected to FLAG IP. Proteins were separated by SDS-PAGE, and analysed by immunoblotting with the indicated antibodies. Data shown are representative of at least 2 independent experiments. The positions of molecular mass markers in kDa are indicated on the left of each immunoblot.

Since porcine and human TBK1 share 96% amino acid identity, and sapovirus proteases have been reported to share cleavage sites, we sought to determine if human sapovirus (HuSaV) NS6 protease also had a similar effect on pTBK1 protein levels. To address this, we cloned wild type HuSaV (GI.2 strain) NS6-7 into a mammalian expression vector. mCherry-tagged HuSaV and PSaV NS6-7 were then overexpressed together with FLAG-tagged pTBK1, or GFP as control. The PSaV NS6-7 was used as a positive control, whereas a condition without a protease was also included. The expression of either PSaV or HuSaV NS6-7 lead to a substantial decrease in TBK1 protein levels (Figure 3C), but not of GFP (Figure 3D).

Thus far, our results revealed a consistent reduction in TBK1 total protein levels, without the detection of an apparent cleavage product. To further investigate the interplay between NS6 and TBK1, and to determine if there is a physical association, we performed immunoprecipitation (IP) assays. Specifically, we overexpressed HA-tagged NS6-7, where NS6 was WT or MUT, or HA-GFP, together with FLAG-tagged pTBK1. A FLAG IP was performed and samples were analysed by immunoblotting. TBK1 was found to interact with NS6-7, but not with GFP (Figure 3E). Consistent with our previous findings (Figure 3A-C), the presence of WT NS6 led to a reduction in TBK1, resulting in less TBK1 being pulled-down compared to the other conditions. Nevertheless, traces of NS6-7 were still identified, indicating the persistence of the NS6-7/TBK1 complex.

### 3.4 TBK1 protein levels are reduced during PSaV infection

To investigate the TBK1 protein levels during infection, we infected LLC-PK1 cells with a high multiplicity of infection (MOI) of PSaV (5 TCID50/cell) and collected samples at different time points post-infection (pi). Viral protein expression was detectable by western blot primarily from 8 h pi onwards, although some VPg precursors were observable at 4 h pi. In line with our previous results (Figure 3), we found that the levels of TBK1 were reduced from 16 h pi (Figure 4A). Quantification of the TBK1 levels observed during PSaV-infected LLC-PK1 cells, revealed that the total TBK1 levels were reduced by at least 50% when compared to mock-infected cells (Figure 4B). To assess whether the decreased expression of TBK1 was cell-type specific, we established PSaV infection in the IPEC-J2 cells, an enterocyte cell line originally derived from the jejunum of a neonatal piglet [23]. We found that the presence of bile acid (GCDCA) was required for efficient infection (Figure 4C), with 300 μM being the optimal concentration (Figure S1). High MOI infection in the IPEC-J2 cells demonstrated that TBK1 levels were reduced (Figure 4D), in agreement with our previous observations (Figure 3 & 4A-B). Quantification of the TBK1 levels in PSaV-infected IPEC-J2 cells showed that the protein levels were decreased by approximately 50% when compare to uninfected cells (Figure 4E), similarly to what we observed in LLC-PK1 cells (Figure 4B).

**Figure 4.**
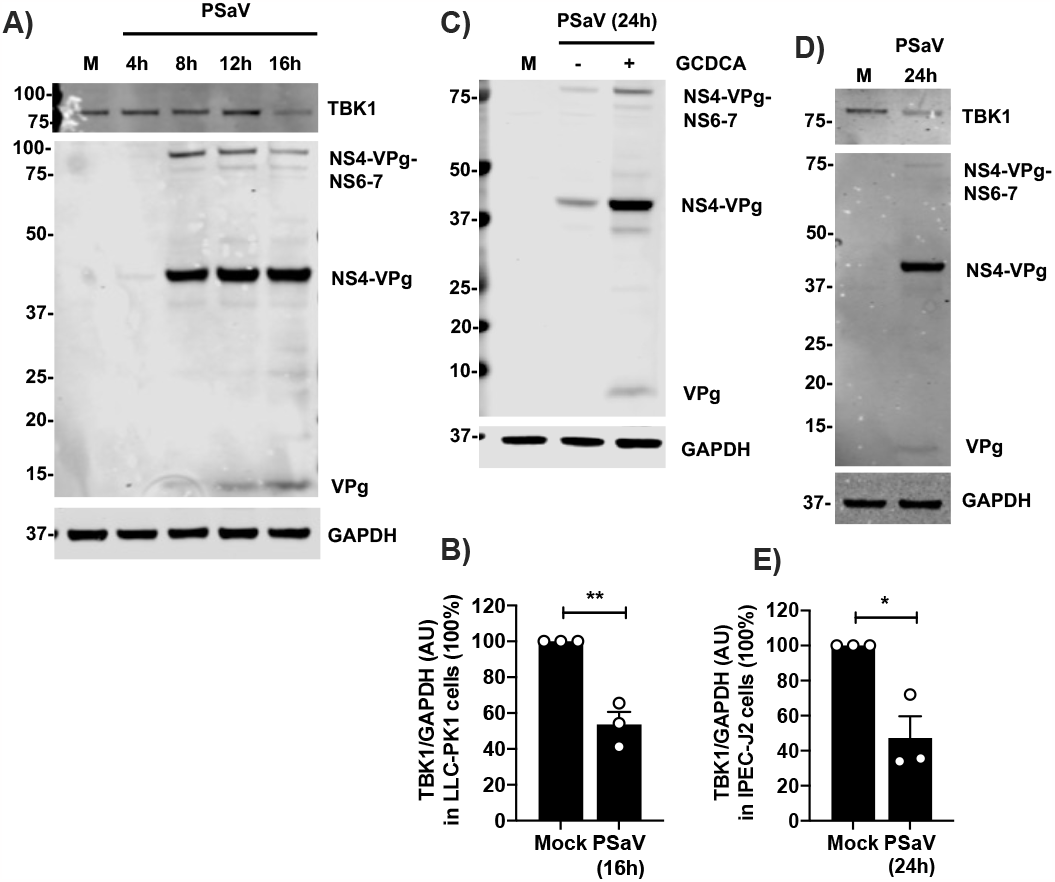
Protein levels of TBK1 are reduced during PSaV infection. A) LLC-PK1 cells mock-infected, or infected with PSaV at 5 TCID50/cell for the indicated h pi. All samples were lysed together, loaded onto a NuPAGE (10 μg/sample) and proteins were separated and analysed by immunoblotting. Immunoblot shown is the representative of 2 independent experiments. B) Arbitrary units (AU) of TBK1 normalised to GAPDH, from mock-, or PSaV-infected LLC-PK1 cells. In all 3 experiments, cells were infected at 5 TCID50/cell and lysed at 16 h pi. C) IPEC-J2 cells mock-infected, or infected with PSaV at 5 TCID50/cell in the presence or absence of GCDCA (300 μM). Cells were lysed at 24 h pi, loaded onto a NuPAGE (10 μg/sample), separated and immunoblotted with the indicated antibodies. Immunoblot shown is the representative of 3 independent experiments. D) IPEC-J2 cells were infected with PSaV at 5 TCID50/cell in the presence of 300 μM GCDCA, and they were lysed at 24 h pi. Proteins were separated by NuPAGE (10 μg/sample), and immunoblotting analysis was followed with the indicated antibodies. Immunoblot shown is the representative of 3 independent experiments. E) AU of TBK1 normalised to GAPDH, of mock- or PSaV-infected IPEC-J2 cells. In all 3 experiments, cells were infected with PSaV at 5 TCID50/cell and cells were lysed at 24 h pi. For all the immunoblots, antibodies used are shown on the right, whereas precursor and mature VPg were detected by an anti-VPg antibody. Positions of molecular mass markers are indicated on the left. Statistical significance was measured by a Student’s *t-*test. **\****p* ≤ 0.05, \*\**p* ≤ 0.01. Error bars represent standard error of the mean.

### 3.5 NS6-mediated reduction of TBK1 levels is proteasomal-dependent

As detailed above, we were unable to identify a TBK1 cleavage product in either infected cells or in cells over-expressing NS6-7. The use of a double-tagged protein also failed to clearly identify a cleavage product (Figure 3B). Therefore, to understand the mechanism of NS6-7 mediated TBK1 degradation, we sought to identify the cellular degradation pathway(s) that could mediate the subsequent turnover of the cleaved TBK1. We treated pTBK1 and NS6-7 co-transfected cells with increasing doses of inhibitors for cellular caspases (zVAD-FMK), lysosomal degradation (NH4Cl) or proteasomal degradation (MG132 and lactacystin). A construct encoding inactive NS6 (MUT) was also included as control. We found that TBK1 levels were unaffected in the presence of NS6 MUT, or when cells were treated with either of the proteasome inhibitors (Figure 5A), suggesting that TBK1 is cleaved by catalytically active NS6 and then targeted for rapid degradation by the proteasome. An additional protein product with a lower molecular mass was detected in lactacystin-treated cells (indicated with blue arrow). Further investigation showed that this anti-TBK1 reactive protein product was present in lactacystin-treated cells independently of the enzymatic activity of NS6 (Figure S2), suggesting that it is an unspecific protein which cross-reacts with the anti-FLAG antibody in the conditions tested.

**Figure 5.**
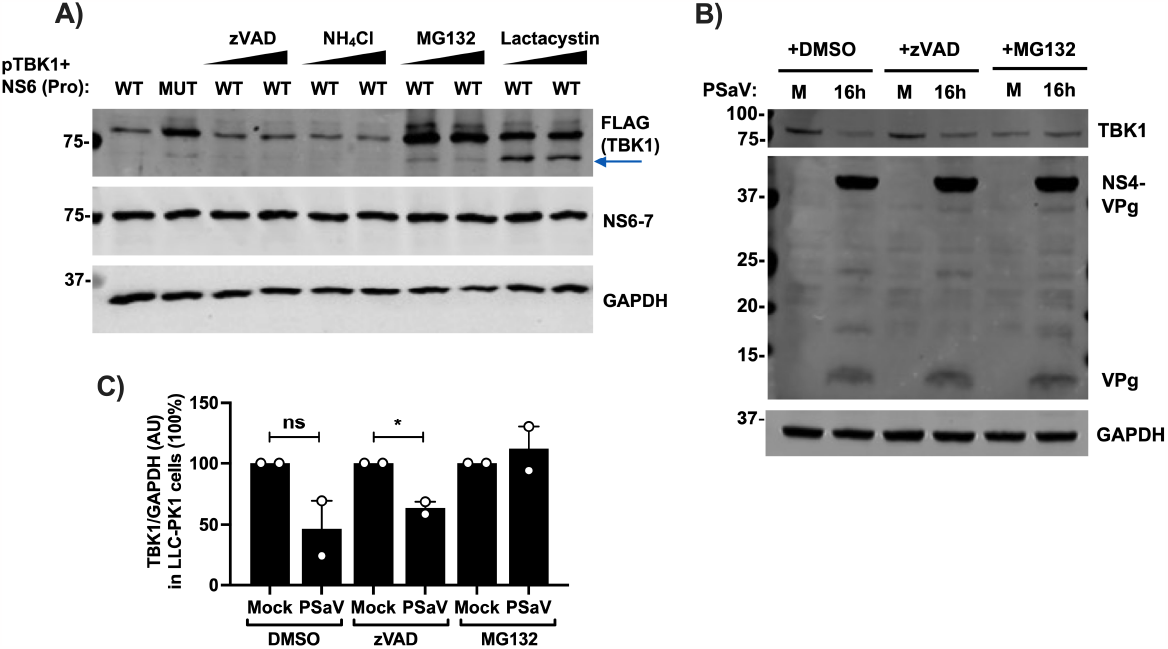
TBK1 is targeted for proteasomal degradation following cleavage by NS6. A) HEK293T cells were co-transfected with 1 ug of HA-NS6-7 where NS6 was WT (Pro^WT^Pol^WT^), or MUT (Pro^MUT^Pol^WT^), and 1 ug of FLAG-pTBK1. Five h post-transfection, cells were treated with zVAD-FMK (10 and 20 μM), or NH_4_Cl (10 and 20 mM), MG132 or lactacystin (both at 5 and 10 μM), or DMSO as control for a total of 24 h. Cells were then lysed, loaded onto an SDS-PAGE (2 μg/sample), separated and analysed by immunoblotting with the indicated antibodies. Positions of molecular mass markers in kDa are indicated on the left. Blue arrow indicates the unspecific band. Immunoblot shown is representative of 3 independent experiments. B) LLC-PK1 cells were infected with PSaV at 5 TCID50/cell. Post inoculum incubation, cells were treated with 20 μM zVAD-FMK or 20 μM MG132, DMSO as control and infection was left for a total of 16 h. Cells were lysed, samples were loaded onto a NuPAGE (10 μg/sample), and proteins were separated and analysed by immunoblotting. Antibodies used are shown on the right, and precursor and mature VPg were detected by an anti-VPg antibody. Positions of molecular mass markers in kDa are indicated on the left. Immunoblot shown is representative of 2 independent experiments. C) AU of TBK1 normalised to GAPDH, of mock- or PSaV-infected LLC-PK1 cells from 2 independent experiments. In both experiments, cells were infected with PSaV at 5 TCID50/cell and cells were treated with the drugs as described in B). Statistical significance was measured by a Student’s *t-*test. ns: non-significant, **\****p* ≤ 0.05. Error bars represent standard error of the mean.

In order to assess whether cleaved TBK1 is also targeted for proteasomal degradation during the course of infection, we infected LLC-PK1 cells at high MOI (5 TCID50/cell) in the presence or absence of zVAD-FMK or MG132. TBK1 levels were reduced in both DMSO- and zVAD-FMK-treated cells upon infection (Figure 5B), which is in accordance with our previous observations (Figure 4 & 5A). In contrast, MG132 treatment, rescued TBK1 (Figure 5B), confirming our findings that TBK1 is targeted for degradation via the proteasome. Quantification of TBK1 levels normalised to GAPDH internal control, further confirmed our observations (Figure 5C).

### 3.6 PSaV infection in TBK1-deficient cells results in higher viral titres

In order to further explore the role of TBK1 in the PSaV life cycle, we sought to examine how lack of TBK1 impacts PSaV replication in cells. For this, we generated TBK1 knockout (KO) LLC-PK1 cells by CRISPR-Cas9-mediated targeting of *Tbk1* exon 3, which is conserved in all porcine TBK1 isoforms that result in the same size protein (Figure 6A). After single cell selection and clonal expansion, cells were confirmed to lack TBK1 by both genomic sequencing (Figure 6A) and immunoblotting (Figure 6B). A clone that lacked TBK1 protein expression and in which the *Tbk1* open reading frame was disrupted by frameshift mutation in both alleles, was selected. In parallel to the KO line, another clonal line was generated by using an empty vector (EV) (see methods for details). The EV line exhibited similar TBK1 expression as the parental line (Figure 6B), and the single clone selection did not influence PSaV replication in these cells (Figure S3). As a further validation, we transfected the EV or KO LLC-PK1 cell lines with poly(I:C) to induce innate immune response, and we measured the *IFN-β* mRNA levels by RT-qPCR. Deletion of TBK1 did not affect *IFN-β* basal levels, but it had a profound effect on *IFN-β* transcription upon stimulation (Figure 6C). We then infected TBK1 WT (EV) or KO cells with PSaV at a low MOI (1 TCID50/cell) and we measured total virus production 24 h pi by TCID50. PSaV titres from TBK1-deficient cells were 0.5-Log10 more than in WT cells, corresponding to 2.6-fold higher virus titres (Figure 6D). Finally, this fold increase on virus yield was consistent and it was observed in 2 independent experiments (Figure 6E). Taken all together, TBK1 plays a crucial role in facilitating type I IFN production in LLC-PK1 cells, and its deletion has a positive effect on PSaV replication and virus yield.

**Figure 6.**
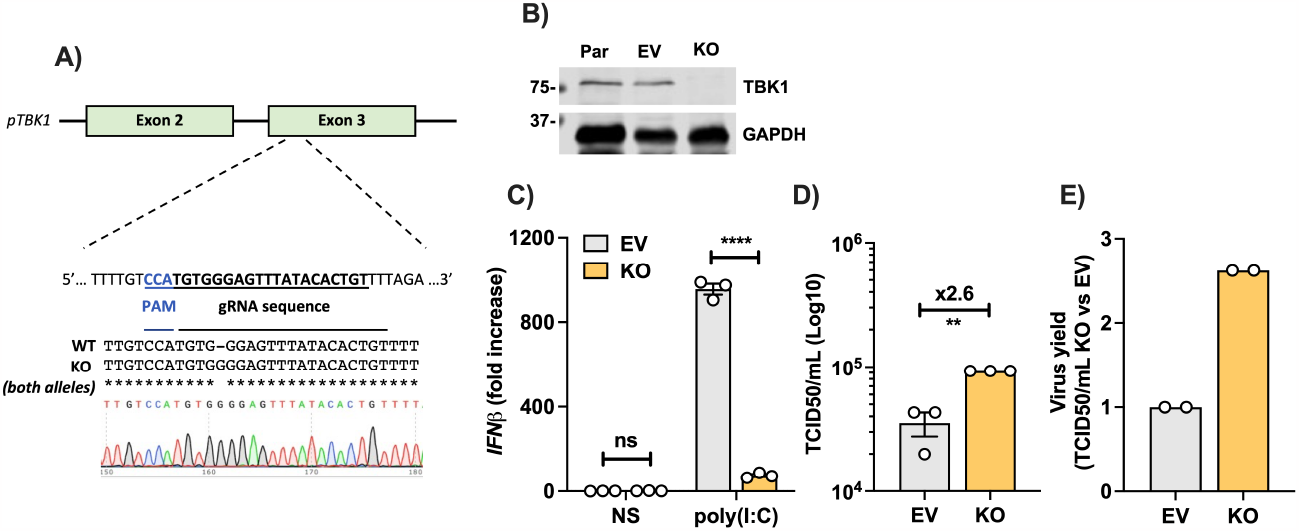
PSaV infection in TBK1-deficient cells results in higher viral titres. A) Schematic of CRISPR-Cas9-mediated knockout strategy targeting porcine *Tbk1* exon 3 and DNA sequences of LLC-PK1 WT and KO cells. Both alleles of KO clone showed the same frameshifting and the chromatogram is included below. B) Immunoblot analysis of LLC-PK1 parental (Par), EV and KO lines for endogenous levels of TBK1 and GAPDH. The positions of molecular mass markers in kDa are shown on the left. C) LLC-PK1 WT (EV) and TBK1 KO cells were transfected with 1 ug/mL poly(I:C) or left untransfected as non-stimulated (NS) control. Six h later, cells were lysed and total RNA was extracted, followed by RT-qPCR. The *IFN-β* levels were compared to *β-actin* levels of each sample, and then everything was compared to EV NS to obtain fold increase. Data shown are representatives of 2 independent experiments done in triplicate. D) TBK1 WT (EV) and KO LLC-PK1 cells were infected with PSaV at 1 TCID50/cell for 24 h. Total virus yield was then measured by TCID50. Data shown are representatives of 2 independent experiments done in triplicate. E) PSaV yield from TBK1 WT (EV) and KO LLC-PK1 cells from 2 independent experiments, represented as fold change. Fold increase was calculated within each experiment. Statistical significance was measured by a Student’s *t-*test. ns: non-significant, \*\**p* ≤ 0.01, \*\*\*\**p* ≤ 0.0001. Error bars represent standard error of the mean.

## 4. Discussion

Viruses constantly evolve to circumvent the host’s IFN response, in order to establish a successful infection. PSaV Cowden strain was the first SaV to successfully propagate *in vitro* [13] and to have a reverse genetic system [14]. It is surprising therefore that our knowledge about SaV-host interactions is currently limited. PSaV is sensitive to type I IFN [18], and thus we sought to understand which mechanisms it utilises to counteract innate immune responses. We found that the 3C-like protease (NS6) of PSaV Cowden strain is an IFN antagonist, by inhibiting the type I IFN production at the level of TBK1. We also showed that TBK1 is targeted by NS6, and for this, the enzymatic activity of NS6 is required. Finally, the role of TBK1 in the IFN response against PSaV was confirmed in immortalised cells deficient for TBK1. PSaV infection in these cells results in a 2.6-fold increase of viral titres.

Positive-sense RNA viruses encode 3C or 3C-like proteins, which are chymotrypsin-like proteases. There is compelling evidence that various viral proteases of RNA viruses function as IFN antagonists by cleaving proteins involved in the IFN response, such as MAVS [34,35], TRIF [35] and NEMO [36]. Recent studies within the *Caliciviridae* family have revealed that the 3C-like proteases of human norovirus and rabbit hemorrhagic disease virus (RHDV) counteract the host’s immune response by cleaving NEMO [31] and MAVS [37], respectively. Whether PSaV NS6 also cleaves these proteins is yet to be determined. However, our results indicate that it also employs alternative mechanisms, as our results demonstrate that it targets TBK1, which is downstream of MAVS.

Despite multiplex experimental approaches, we did not detect any apparent NS6 cleavage product of TBK1. Notably, porcine picornavirus and poliovirus 3C proteases have been found to induce degradation of cellular targets without without the appearance of readily detectable cleavage products [38,39]. One plausible explanation for our results is that TBK1 may undergo cleavage at a proximal site, leading to rapid instability. In contrast to the porcine picornavirus and poliovirus studies detailed above, here we have demonstrated that TBK1 is degraded via the proteasome, as MG132 and lactacystin treatments restored both ectopic and endogenous levels of TBK1. The ability of viruses to target TBK1 is not unique to PSaV, as there is evidence of other viruses to either cleave TBK1 [40], or induce its degradation [41], as a means to impair the host’s IFN response.

Our findings indicate that TBK1 is targeted during late stages of infection, and there are several potential reasons for this observation. Firstly, it is plausible that NS6 does not co-localise with TBK1 during earlier stages of the virus life cycle, or that TBK1 activation and thus its adoption of a different structural conformation [42] are necessary for their interaction to occur. Moreover, TBK1 may hold important pro-viral roles earlier during infection. A recent study demonstrated that PSaV induces RIPK1-dependent necroptosis, which promotes virus replication [43]. TBK1, along with IKK-ε and other IKKs, phosphorylates RIPK1, negatively regulating its kinase activity and ultimately promoting cell survival [44]. Therefore, it is possible that NS6 targets TBK1 at a specific time pi to finely regulate the phosphorylation and activity of RIPK1. It should be noted that although TBK1 was deleted from LLC-PK1 cells, we do not anticipate a decrease in the expression levels of IKKs. This suggests that the kinase activity of RIPK1 may still be regulated to some extent. Finally, while our study has demonstrated that TBK1 is targeted during late stages of PSaV infection, it is important to acknowledge that the virus may exhibit additional mechanisms to circumvent the innate immune response earlier in the infection process. These mechanisms, have not been examined and are the subject of future investigations.

While LLC-PK1 cells have been widely used as a model for propagating PSaV, they are kidney cells and likely do not represent the natural target for PSaV. In this study, we successfully established PSaV infection in IPEC-J2 cells by supplementing the culture medium with bile acids. Specifically, we tested Glycochenodeoxycholic acid (GCDCA), although it is possible that other bile acids may enhance PSaV replication to a greater extent. IPEC-J2 cells are derived from the jejunum of neonatal piglets [23], which represents one of the natural hosts for PSaV [15]. These cells exhibit striking similarities to primary intestinal epithelial cells, making them a valuable tool for investigating the interactions between the porcine gut epithelium and enteric viruses and bacteria [45,46]. To the best of our knowledge, this study represents the first instance of PSaV infection in IPEC-J2 cells. Thus, we have provided a new, relevant tool for further investigations of PSaV replication cycle, pathogenesis, and host response in the context of the porcine intestinal environment.

Our study provides the first evidence of a SaV protein to act as innate immune antagonist. Our discovery that HuSaV NS6 also targets porcine TBK1, which shares over 96% aa identity with human TBK1, could be significant for HuSaV research. Furthermore, considering the genetic similarity between some PSaVs and HuSaVs [16], our findings may suggest a similar mechanism employed by HuSaV to circumvent the type I IFN response. Recent advancements in propagating HuSaV in diverse cell lines [12] and human intestinal enteroids [10], have expanded the opportunities for studying HuSaV. Thus, our findings offer a promising avenue to deepen our understanding of the mechanisms of HuSaV infection and interactions with its host.

## Supplementary Materials

Figure S1: 300 μM is the optimal concentration of GCDCA for PSaV infection in IPEC-J2 cells.; Figure S2: Presence of a non-specific band in lactacystin-treated cells, independently of NS6’s enzymatic activity.; Figure S3: Clonal selectivity of LLC-PK1 EV cells does not affect PSaV replication.

## Author Contributions

Conceptualization: I.G., M.H. and I.G.G.; Validation: I.G. and M.H.; Formal analysis: I.G. and M.H.; Investigation: I.G., M.H., A.S.J., F.S. and E.E.; Resources: I.G.G.; Data curation: I.G. and M.H.; Writing—original draft preparation: I.G.; Writing—review and editing: I.G., M.H., A.S.J., E.E., F.S., K.O.C., and I.G.G.; Visualization: I.G.; Supervision: I.G.G.; Funding acquisition: I.G.G. All authors have read and agreed to the published version of the manuscript.

## Funding

This research was funded by the Wellcome Trust, grant number 207498/Z/17/Z.

## Data Availability Statement

Data for this submission have been deposited at Figshare, DOI 10.6084/m9.figshare.24757419.

## Acknowledgments

We thank Prof. Geoffrey L. Smith (University of Oxford) for providing plasmids for this study.

## Conflicts of Interest

Authors disclose no conflicts of interest.

## Notes

### Competing Interest Statement

The authors have declared no competing interest.

## References

1. Li, J.; Zhang, W.; Cui, L.; Shen, Q.; Hua, X. Metagenomic identification, genetic characterization and genotyping of porcine sapoviruses. Infection, Genetics and Evolution 2018, 62, 244–252, doi:10.1016/j.meegid.2018.04.034.

2. Yinda, C.K.; Conceição-Neto, N.; Zeller, M.; Heylen, E.; Maes, P.; Ghogomu, S.M.; Van Ranst, M.; Matthijnssens, J. Novel highly divergent sapoviruses detected by metagenomics analysis in straw-colored fruit bats in Cameroon. Emerging Microbes & Infections 2017, 6, 1–7, doi:10.1038/emi.2017.20.

3. Oka, T.; Wang, Q.; Katayama, K.; Saif Linda, J. Comprehensive Review of Human Sapoviruses. Clinical Microbiology Reviews 2015, 28, 32–53, doi:10.1128/cmr.00011-14.

4. Oka, T.; Katayama, K.; Ogawa, S.; Hansman, G.S.; Kageyama, T.; Ushijima, H.; Miyamura, T.; Takeda, N. Proteolytic Processing of Sapovirus ORF1 Polyprotein. Journal of Virology 2005, 79, 7283–7290, doi:10.1128/jvi.79.12.7283-7290.2005.

5. Li, T.-C.; Kataoka, M.; Doan, Y.H.; Saito, H.; Takagi, H.; Muramatsu, M.; Oka, T. Characterization of a Human Sapovirus Genotype GII.3 Strain Generated by a Reverse Genetics System: VP2 Is a Minor Structural Protein of the Virion. Viruses 2022, 14, 1649, doi:10.3390/v14081649.

6. Oka, T.; Katayama, K.; Ogawa, S.; Hansman, G.S.; Kageyama, T.; Ushijima, H.; Miyamura, T.; Takeda, N. Proteolytic processing of sapovirus ORF1 polyprotein. J Virol 2005, 79, 7283–7290, doi:10.1128/jvi.79.12.7283-7290.2005.

7. Sosnovtseva, S.A.; Sosnovtsev, S.V.; Green, K.Y. Mapping of the feline calicivirus proteinase responsible for autocatalytic processing of the nonstructural polyprotein and identification of a stable proteinase-polymerase precursor protein. J Virol 1999, 73, 6626–6633, doi:10.1128/jvi.73.8.6626-6633.1999.

8. Chiba, S.; Sakuma, Y.; Kogasaka, R.; Akihara, M.; Horino, K.; Nakao, T.; Fukui, S. An outbreak of gastroenteritis associated with calicivirus in an infant home. Journal of Medical Virology 1979, 4, 249–254, doi:10.1002/jmv.1890040402.

9. Madeley, C.R.; Cosgrove, B.P. CALICIVIRUSES IN MAN. The Lancet 1976, 307, 199–200, doi:10.1016/S0140-6736(76)91309-X.

10. Euller-Nicolas, G.; Le Mennec, C.; Schaeffer, J.; Zeng, X.-L.; Ettayebi, K.; Atmar Robert, L.; Le Guyader Françoise, S.; Estes Mary, K.; Desdouits, M. Human Sapovirus Replication in Human Intestinal Enteroids. Journal of Virology 2023, 97, e00383–00323, doi:10.1128/jvi.00383-23.

11. Matsumoto, N.; Kurokawa, S.; Tamiya, S.; Nakamura, Y.; Sakon, N.; Okitsu, S.; Ushijima, H.; Yuki, Y.; Kiyono, H.; Sato, S. Replication of Human Sapovirus in Human-Induced Pluripotent Stem Cell-Derived Intestinal Epithelial Cells. Viruses 2023, 15, doi:10.3390/v15091929.

12. Takagi, H.; Oka, T.; Shimoike, T.; Saito, H.; Kobayashi, T.; Takahashi, T.; Tatsumi, C.; Kataoka, M.; Wang, Q.; Saif, L.J.; et al. Human sapovirus propagation in human cell lines supplemented with bile acids. Proceedings of the National Academy of Sciences 2020, 117, 32078–32085, doi:10.1073/pnas.2007310117.

13. Chang, K.O.; Sosnovtsev, S.V.; Belliot, G.; Kim, Y.; Saif, L.J.; Green, K.Y. Bile acids are essential for porcine enteric calicivirus replication in association with down-regulation of signal transducer and activator of transcription 1. Proc Natl Acad Sci U S A 2004, 101, 8733–8738, doi:10.1073/pnas.0401126101.

14. Chang, K.O.; Sosnovtsev, S.V.; Belliot, G.; Wang, Q.; Saif, L.J.; Green, K.Y. Reverse genetics system for porcine enteric calicivirus, a prototype sapovirus in the Caliciviridae. J Virol 2005, 79, 1409–1416, doi:10.1128/jvi.79.3.1409-1416.2005.

15. Nagai, M.; Wang, Q.; Oka, T.; Saif, L.J. Porcine sapoviruses: Pathogenesis, epidemiology, genetic diversity, and diagnosis. Virus Res 2020, 286, 198025, doi:10.1016/j.virusres.2020.198025.

16. Martella, V.; Lorusso, E.; Banyai, K.; Decaro, N.; Corrente, M.; Elia, G.; Cavalli, A.; Radogna, A.; Costantini, V.; Saif, L.J.; et al. Identification of a porcine calicivirus related genetically to human sapoviruses. J Clin Microbiol 2008, 46, 1907–1913, doi:10.1128/jcm.00341-08.

17. Jahun, A.S.; Goodfellow, I.G. Interferon responses to norovirus infections: current and future perspectives. J Gen Virol 2021, 102, doi:10.1099/jgv.0.001660.

18. Hosmillo, M.; Sorgeloos, F.; Hiraide, R.; Lu, J.; Goodfellow, I.; Cho, K.O. Porcine sapovirus replication is restricted by the type I interferon response in cell culture. J Gen Virol 2015, 96, 74–84, doi:10.1099/vir.0.071365-0.

19. Rehwinkel, J.; Gack, M.U. RIG-I-like receptors: their regulation and roles in RNA sensing. 2020, doi:10.1038/s41577-020-0288-3.

20. Carty, M.; Guy, C.; Bowie, A.G. Detection of Viral Infections by Innate Immunity. Biochemical Pharmacology 2021, 183, 114316, doi:10.1016/j.bcp.2020.114316.

21. Fitzgerald, K.A.; McWhirter, S.M.; Faia, K.L.; Rowe, D.C.; Latz, E.; Golenbock, D.T.; Coyle, A.J.; Liao, S.-M.; Maniatis, T. IKKε and TBK1 are essential components of the IRF3 signaling pathway. Nature Immunology 2003, 4, 491–496, doi:10.1038/ni921.

22. Smale, S.T. Selective Transcription in Response to an Inflammatory Stimulus. Cell 2010, 140, 833–844, doi:10.1016/j.cell.2010.01.037.

23. Berschneider, H. Development of normal cultured small intestinal epithelial cell lines which transport Na and Cl. Abstract of the Annual Meeting of the American Gastroenterological Association. In Proceedings of the Digestive Disease Week and the 90th annual meeting of the American Gastroenterological Association. Elsevier. Washington DC, 1989.

24. Geens, M.M.; Niewold, T.A. Optimizing culture conditions of a porcine epithelial cell line IPEC-J2 through a histological and physiological characterization. Cytotechnology 2011, 63, 415–423, doi:10.1007/s10616-011-9362-9.

25. Odon, V.; Georgana, I.; Holley, J.; Morata, J.; Maluquer de Motes, C. Novel Class of Viral Ankyrin Proteins Targeting the Host E3 Ubiquitin Ligase Cullin-2. J Virol 2018, 92, doi:10.1128/jvi.01374-18.

26. Lin, R.; Heylbroeck, C.; Pitha, P.M.; Hiscott, J. Virus-Dependent Phosphorylation of the IRF-3 Transcription Factor Regulates Nuclear Translocation, Transactivation Potential, and Proteasome-Mediated Degradation. Molecular and Cellular Biology 1998, 18, 2986–2996, doi:10.1128/MCB.18.5.2986.

27. Ranjith-Kumar, C.T.; Wen, Y.; Baxter, N.; Bhardwaj, K.; Cheng Kao, C. A Cell-Based Assay for RNA Synthesis by the HCV Polymerase Reveals New Insights on Mechanism of Polymerase Inhibitors and Modulation by NS5A. PLOS ONE 2011, 6, e22575, doi:10.1371/journal.pone.0022575.

28. Ran, F.A.; Hsu, P.D.; Wright, J.; Agarwala, V.; Scott, D.A.; Zhang, F. Genome engineering using the CRISPR-Cas9 system. Nature Protocols 2013, 8, 2281–2308, doi:10.1038/nprot.2013.143.

29. Reed, L.J.; Muench, H. A SIMPLE METHOD OF ESTIMATING FIFTY PER CENT ENDPOINTS12. American Journal of Epidemiology 1938, 27, 493–497, doi:10.1093/oxfordjournals.aje.a118408.

30. Lei, J.; Hilgenfeld, R. RNA-virus proteases counteracting host innate immunity. FEBS Lett 2017, 591, 3190–3210, doi:10.1002/1873-3468.12827.

31. Zhang, H.; Jiang, P.; Chen, Z.; Wang, D.; Zhou, Y.; Zhu, X.; Xiao, S.; Fang, L. Norovirus 3C-Like protease antagonizes interferon-β production by cleaving NEMO. Virology 2022, 571, 12–20, doi:10.1016/j.virol.2022.04.004.

32. Hornung, V.; Ellegast, J.; Kim, S.; Brzózka, K.; Jung, A.; Kato, H.; Poeck, H.; Akira, S.; Conzelmann, K.-K.; Schlee, M.; et al. 5’-Triphosphate RNA Is the Ligand for RIG-I. Science 2006, 314, 994–997, doi:10.1126/science.1132505.

33. Pichlmair, A.; Schulz, O.; Tan, C.P.; Näslund, T.I.; Liljeström, P.; Weber, F.; Reis e Sousa, C. RIG-I-Mediated Antiviral Responses to Single-Stranded RNA Bearing 5’-Phosphates. Science 2006, 314, 997–1001, doi:10.1126/science.1132998.

34. Yang, Y.; Liang, Y.; Qu, L.; Chen, Z.; Yi, M.; Li, K.; Lemon, S.M. Disruption of innate immunity due to mitochondrial targeting of a picornaviral protease precursor. Proceedings of the National Academy of Sciences 2007, 104, 7253–7258, doi:10.1073/pnas.0611506104.

35. Mukherjee, A.; Morosky, S.A.; Delorme-Axford, E.; Dybdahl-Sissoko, N.; Oberste, M.S.; Wang, T.; Coyne, C.B. The Coxsackievirus B 3Cpro Protease Cleaves MAVS and TRIF to Attenuate Host Type I Interferon and Apoptotic Signaling. PLOS Pathogens 2011, 7, e1001311, doi:10.1371/journal.ppat.1001311.

36. Wang, D.; Fang, L.; Li, K.; Zhong, H.; Fan, J.; Ouyang, C.; Zhang, H.; Duan, E.; Luo, R.; Zhang, Z.; et al. Foot-and-Mouth Disease Virus 3C Protease Cleaves NEMO To Impair Innate Immune Signaling. Journal of Virology 2012, 86, 9311–9322, doi:10.1128/jvi.00722-12.

37. Men, Y.; Wang, Y.; Wang, H.; Zhang, M.; Liu, J.; Chen, Y.; Han, X.; Chen, R.; Chen, Q.; Hu, A. RHDV 3C protein antagonizes type I interferon signaling by cleaving interferon promoter stimulated 1 protein. Virus Genes 2023, 59, 215–222, doi:10.1007/s11262-022-01958-w.

38. Weidman, M.K.; Yalamanchili, P.; Ng, B.; Tsai, W.; Dasgupta, A. Poliovirus 3C Protease-Mediated Degradation of Transcriptional Activator p53 Requires a Cellular Activity. Virology 2001, 291, 260–271, doi:10.1006/viro.2001.1215.

39. Wang, C.; Feng, H.; Zhang, X.; Li, K.; Yang, F.; Cao, W.; Liu, H.; Gao, L.; Xue, Z.; Liu, X.; et al. Porcine Picornavirus 3C Protease Degrades PRDX6 to Impair PRDX6-mediated Antiviral Function. Virologica Sinica 2021, 36, 948–957, doi:10.1007/s12250-021-00352-4.

40. Jeremiah, S.S.; Miyakawa, K.; Matsunaga, S.; Nishi, M.; Kudoh, A.; Takaoka, A.; Sawasaki, T.; Ryo, A. Cleavage of TANK-Binding Kinase 1 by HIV-1 Protease Triggers Viral Innate Immune Evasion. Frontiers in Microbiology 2021, 12, doi:10.3389/fmicb.2021.643407.

41. Su, W.; Lin, X.-T.; Zhao, S.; Zheng, X.-Q.; Zhou, Y.-Q.; Xiao, L.-L.; Chen, H.; Zhang, Z.-Y.; Zhang, L.-J.; Wu, X.-X. Tripartite motif-containing protein 46 accelerates influenza A H7N9 virus infection by promoting K48-linked ubiquitination of TBK1. Virology Journal 2022, 19, 176, doi:10.1186/s12985-022-01907-x.

42. Larabi, A.; Devos Juliette M.; Ng, S.-L.; Nanao Max H.; Round, A.; Maniatis, T.; Panne, D. Crystal Structure and Mechanism of Activation of TANK-Binding Kinase 1. Cell Reports 2013, 3, 734–746, doi:10.1016/j.celrep.2013.01.034.

43. Sharif, M.; Baek, Y.-B.; Nguyen, T.H.; Soliman, M.; Cho, K.-O. Porcine sapovirus-induced RIPK1-dependent necroptosis is proviral in LLC-PK cells. PLOS ONE 2023, 18, e0279843, doi:10.1371/journal.pone.0279843.

44. Delanghe, T.; Dondelinger, Y.; Bertrand, M.J.M. RIPK1 Kinase-Dependent Death: A Symphony of Phosphorylation Events. Trends in Cell Biology 2020, 30, 189–200, doi:10.1016/j.tcb.2019.12.009.

45. Schierack, P.; Nordhoff, M.; Pollmann, M.; Weyrauch, K.D.; Amasheh, S.; Lodemann, U.; Jores, J.; Tachu, B.; Kleta, S.; Blikslager, A.; et al. Characterization of a porcine intestinal epithelial cell line for in vitro studies of microbial pathogenesis in swine. Histochemistry and Cell Biology 2006, 125, 293–305, doi:10.1007/s00418-005-0067-z.

46. Brosnahan, A.J.; Brown, D.R. Porcine IPEC-J2 intestinal epithelial cells in microbiological investigations. Veterinary Microbiology 2012, 156, 229–237, doi:10.1016/j.vetmic.2011.10.017.

