## Supplementary figures and images for "Porcine sapovirus protease controls the innate immune response and targets TBK1"

### Supplementary figures

S1

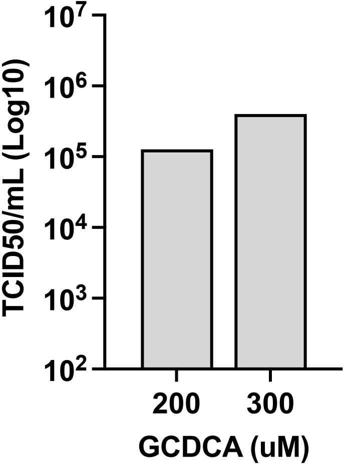

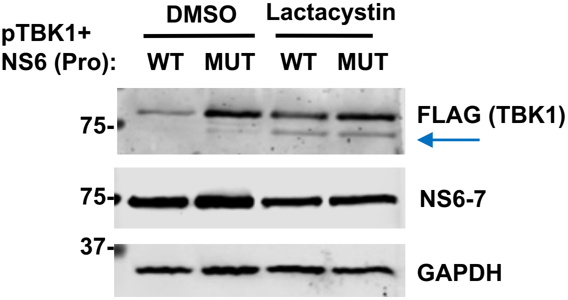

S3

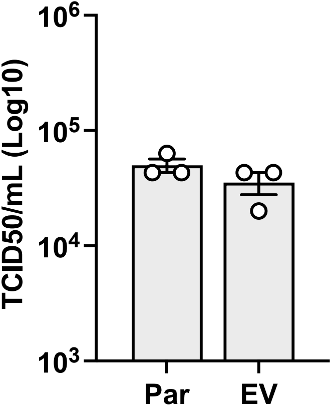
